# MYCN driven oncogenesis involves cooperation with WDR5 to activate canonical MYC targets and G9a to repress differentiation genes

**DOI:** 10.1101/2023.07.11.548643

**Authors:** Zhihui Liu, Xiyuan Zhang, Man Xu, Jason J. Hong, Amanda Ciardiello, Haiyan Lei, Jack F. Shern, Carol J. Thiele

**Affiliations:** Pediatric Oncology Branch, National Cancer Institute, Bethesda, MD, USA

## Abstract

MYCN activates canonical MYC targets involved in ribosome biogenesis, protein synthesis and represses neuronal differentiation genes to drive oncogenesis in neuroblastoma (NB). How MYCN orchestrates global gene expression remains incompletely understood. Our study finds that MYCN binds promoters to up-regulate canonical MYC targets but binds to both enhancers and promoters to repress differentiation genes. MYCN-binding also increases H3K4me3 and H3K27ac on canonical MYC target promoters and decreases H3K27ac on neuronal differentiation gene enhancers and promoters. WDR5 is needed to facilitate MYCN promoter binding to activate canonical MYC target genes, whereas MYCN recruits G9a to enhancers to repress neuronal differentiation genes. Targeting both MYCN’s active and repressive transcriptional activities using both WDR5 and G9a inhibitors synergistically suppresses NB growth. We demonstrate that MYCN cooperates with WDR5 and G9a to orchestrate global gene transcription. The targeting of both these cofactors is a novel therapeutic strategy to indirectly target the oncogenic activity of *MYCN*.

## Introduction

The deregulation of *MYC* family oncogenes including *c-MYC*, *MYCN* and *MYCL* occurs in most cancers and frequently marks those associated with poor prognosis (1–5). *MYCN* is implicated in many pediatric embryonal tumors such as neuroblastoma (NB), rhabdomyosarcoma, medulloblastoma and more recently in therapy-resistant adult cancers including subtypes of breast cancer and prostate cancers (2,5). *MYCN* encodes a basic helix-loop-helix-leucine zipper transcription factor (TF) named N-Myc or MYCN and exhibits high-structural homology with c-MYC (2). Early studies indicated that *MYCN* overexpression in NB cells leads to a transcriptome enriched in canonical *MYC* target genes including genes involved in ribosome biogenesis and protein synthesis (6,7). Later, the identification of a functional MYCN signature gene set in one NB cell line indicated that MYCN suppresses genes associated with neuronal differentiation (8). However, the molecular mechanisms by which MYCN orchestrates these changes in global gene expression at a genome-wide level remain unclear.

Most eukaryotic TFs act by recruiting coactivators, or corepressors, which include chromatin remodeling complexes and covalent histone-modifying complexes (9). Only a handful of coactivators and corepressors of MYCN have been experimentally demonstrated to mediate its transcriptional activity, and in these studies only a few target genes have been assessed (5,10–15). However, the cooperation between MYCN and its cofactors has not been systematically investigated on a genome-wide level. *MYCN* is a *bona-fide* oncogenic driver in NB (5,16–18) and it is known that the silencing of *MYCN* results in a decrease in cell proliferation and induction of cell differentiation in NB cells (19,20). Thus, targeting MYCN has been thought of as a therapeutic aim, but direct MYCN targeting remains challenging due to its structural flexibility. There is the potential for therapeutic strategies aimed at indirect MYCN targeting since cofactors with enzymatic activity are often druggable, but this requires a genome-wide understanding of the critical cofactors by which MYCN regulates global gene expression.

To investigate how MYCN globally regulates gene transcription, we combined a protein interactome assay and genome-wide approaches including RNA sequencing (RNA-seq) and chromatin immunoprecipitation followed by DNA sequencing (ChIP-seq). Our study demonstrates that WDR5 assists MYCN to bind promoters to activate canonical MYC targets whereas MYCN recruits G9a to enhancers to repress neuronal differentiation genes in NB. Importantly, the simultaneous targeting of both WDR5 and G9a-regulated transcriptional activities represents a more effective approach to indirectly target MYCN and its oncogenic program.

## Materials and Methods

### Cell culture

Human neuroblastoma (NB) cell lines IMR32, SK-N-BE(2)C (BE(2)C), SMS-KCNR (KCNR), LAN5 and SHEP were obtained from the cell line bank of the Pediatric Oncology Branch of the National Cancer Institute and have been genetically verified. Human rhabdomyosarcoma cell line RH4 was from Dr. Javed Khan’s lab of the Genetic Branch of the National Cancer Institute. All the NB cell lines, and rhabdomyosarcoma cell line were maintained in RPMI-1640 medium. All the cell culture medium was supplemented with 10% fetal calf serum (FBS), 100 µg/mL streptomycin, 100 U/mL penicillin, and 2 mM L-glutamine. Cells were grown at 37℃ with 5% CO2. All cell lines were frequently assayed for *Mycoplasma* using MycoAlert Kit (Lonza) to ensure they were free of *Mycoplasma* contamination. The cell lines used were within 12 passages after thawing.

### Stable clones

HA tagged MYCN (HA-MYCN) construct was generously provided by Dr. Wei Gu’s lab(21). MYCN coding region was PCR amplified from HA-MYCN construct and cloned into the doxycycline inducible pLVX-pTetOne-puro vector (Takara Bio) using In-Fusion HD kit (Takara Bio) following the manufacturer’s manual. SHEP cells were infected with lentiviral particles generated using the pLVX-TetOne-Puro-MYCN vector, followed by puromycin (0.65 µg/ml) selection. The transduced stable cell line was named as SHEPtetMYCN. MYCN expression in SHEPtetMYCN could be induced with 0.25 µg/ml Dox treatment.

### Transient transfection

siRNA control (AllStars Negative Control siRNA, Catalog No. 1027281) and siRNAs targeting different genes (Hs_MYCN_2, Catalog No. SI00076293; Hs_MYCN_4, Catalog No. SI00076307; Hs_G9a_3, Catalog No. SI00091203; Hs_WDR5_3, Catalog No. SI00118916; Hs_WDR5_4, Catalog No. SI00118923) were purchased from Qiagen or Santa Cruz Biotechnology. siRNAs were transiently transfected into NB cells using Nucleofector electroporation (Lonza): solution L and program C-005 for IMR32; solution V and program A-030 for the rest NB cell lines; solution R and program T-016 for RH4 cell line.

### Cell growth and neurite extension assay

To evaluate cell proliferation, NB cells were plated in 96-well plates and the growth kinetics were monitored in IncuCyte ZOOM or FLR (Essen BioScience) using the integrated confluence algorithm as a surrogate for cell number. Cell neurite length was measured using Essen IncuCyte ZOOM neurite analysis software.

### Monitoring of synergistic effects of drug combinations

The therapeutic effect of WDR5 inhibitor (WDR5i) OICR-9429 (Selleckchem, Catalog No. S7833) and G9a inhibitor (G9ai) UNC0642 (Selleckchem, Catlog No. S7230) in MYCN-Amp NB cell lines IMR32, KCNR and IMR5 was determined in a checkerboard fashion. Cell lines were seeded in two 96-well plates and incubated overnight. Each combination dose had 2 replications. The next day, cell lines were treated with different dose combinations of OICR-9429 and UNC0642. Control cells were treated with DMSO. Each plate has its control cells. Cell viability was determined after 72 h using the CellTiter-Glo® luminescent assay (Promega, catalog number G9242). Cell viability of DMSO-treated cells was set to 100%. Results were graphed with GraphPad Prism (RRID:SCR_002798). IncuCyte® assay was used for testing the impact of synergistic effects of drug combinations on NB cell growth in realtime. Representative data from biological replicates were shown in this study. SynergyFinder (RRID:SCR_019318) online tool (22) was used to study the synergistic effect of the combination treatment of NB cells *in vitro*.

### Soft agar assay

To assess the effects of overexpression of *MYCN* in SHEP cells on anchorage independent cell growth, 1 x 10^4^ SHEPtetMYCN cells were cultured in 0.7% top agarose in media on a layer of 1.4% bottom agar/media to prevent the adhesion of cells to the culture plates. Medium was changed twice a week with or without 0.5 μg/mL Dox, and visible colonies were observed after 2-4 weeks of culture. The number of colonies was counted after crystal violet staining.

### Protein isolation, western blotting analysis, and co-immunoprecipitation

For assessment of protein levels, cells were lysed using RIPA buffer, and 10 µg of total protein was separated and electroblotted. Protein bands probed with diluted primary antibodies (Table S8) were detected using a goat anti-rabbit or mouse IgG-HRP conjugated secondary antibody (Santa Cruz Biotechnology) and visualized using enhanced chemiluminescence (Amersham Biosciences).

To identify the interactome of the endogenous MYCN, co-immunoprecipitation (co-IP) was performed as previously described with slight modification (23). IMR32 cells were solubilized for 30 min in cold lysis buffer (50 mM pH 7.5 Tris-HCl, 137 mM NaCl, 1mM DTT, 1mM EDTA, 0.5% Triton X-100) supplemented with protease and phosphatase inhibitors (Halt protease and phosphatase inhibitor, Thermo), by shaking at 4°C. Two different MYCN antibodies (antibody 1, Santa Cruz, sc-53993; antibody 2, Abcam, ab16898) (4 µg), or normal IgG (4 µg) was incubated with 50 µl Dynabeads M-280 sheep anti-mouse IgG magnetic beads (Thermo Fisher Scientific Cat# 11201D, RRID:AB_2783640) in 200ul wash buffer (50 mM pH 7.5 Tris-HCl, 137 mM NaCl, 1mM EDTA, 0.5% Triton X-100) overnight with rotation at 4 ℃. The clear cell lysate (20 mg) was incubated with the Magnetic Beads coupled with MYCN antibody 1, MYCN antibody 2, or IgG control in total 4.5 ml lysis buffer and agitated at 4°C for 4 h. Subsequently the beads were washed 5 times with washing buffer. The co-IP products were eluted by incubating with 25 µl 1x LDS-PAGE sample buffer supplemented with 10% β-mercaptoethanol and boiling for 5 min. After staining with SimplyBlue Safe Stain reagents, the differentially pulled-down bands were sequenced using mass spectrometry (mass-spec) (NCI-Frederick protein analysis core facility). To validate the mass-spec result, co-IP and western blot were performed, and in this validation experiment, Benzonase (500 U/ml), Mg2+ (2mM) will be added to the co-IP reaction as indicated. Benzonase is a nuclease that digests both DNA and RNA. Of note, for the mass-spec assay, we did not add Benzonase since we aim to identify most of MYCN interactors including those weak interactions that occur when the complex bind to nucleic acids. Primary antibodies of MYCN, G9a and WDR5 (Table S8) were used to detect the protein-protein interaction.

### RNA-seq

Total RNA was isolated from neuroblastoma or rhabdomyosarcoma cells that have been transiently transfected with different siRNAs or siCtrl for 48 h or 72 h and subjected to RNA-seq analysis as previously described (24). Total RNA was extracted using the RNeasy Plus Mini Kit (Qiagen Inc.) according to the manufacturer’s instructions. TruSeq® Stranded Total RNA LT Library Prep Kit or TruSeq Stranded mRNA Library Prep kit (Illumina, San Diego, CA, USA) was used for preparing Strand-specific whole transcriptome sequencing libraries by following the manufacturer’s procedure. The Fastq files with paired-end reads were processed using Partek Flow. The raw reads are aligned using STAR (RRID:SCR_004463) and the aligned reads are quantified to the annotation model through Partek E/M. The normalization method used here is counts per million (CPM) through Partek Flow. The statistic analysis of normalized counts used GSA or ANOVA. To get T-scores, the normalized counts acquired from Partek Flow are exported and further analyzed using Parteck Genomics Suite v7.17. Statistical results of differentially expressed genes from Partek Flow were analyzed using QIAGEN’s Ingenuity® Pathway Analysis (IPA®, QIAGEN) and gene set enrichment analysis (GSEA). By default, the false discovery rate (FDR) less than 0.25 is significant in GSEA.

### ChIP-seq

ChIP-seq was performed using the ChIP-IT High Sensitivity kit (Active Motif, cat. 53040) as described previously (24). Briefly, formaldehyde (1%, 13 minutes) fixed cells were sheared to achieve chromatin fragmented to a range of 200-700 bp using an Active Motif EpiShear Probe Sonicator. IMR32 cells that have been transiently transfected with different siRNAs that target different genes or negative control siRNA (siCtrl) for 72 h were used for ChIP-seq. IMR32 cells were sonicated at 25% amplitude, pulse for 20 seconds on and 30 seconds off for a total sonication “on” time of 16 minutes. Sheared chromatin samples were immunoprecipitated overnight at 4 °C with antibodies targeting MYCN, G9a, WDR5, H3K27ac, H3K4me1, H3K4me3 and H3K27me3 (Table S8). To compare the colocalization between MYCN and other proteins on the genome, ChIP-seq data from negative control siRNA transfected IMR32 cells were used. ChIP-seq DNA libraries were prepared by Frederick National Laboratory for Cancer Research sequencing facility. Libraries were multiplexed and sequenced using TruSeq ChIP Samples Prep Kit (75 cycles), cat. # IP-2-2-1012/1024 on an Illumina NextSeq machine. 25,000,000-30,000,000 unique reads were generated per sample.

### ChIP-seq data processing

Previously published ChIP-seq datasets are down-loaded for this study, which includes ChIP-seq datasets of MYCN that were generated in BE(2)C cells (GSE94822). As described previously (24), for the home generated ChIP-seq data, ChIP enriched DNA reads were mapped to reference the human genome (version hg19) using BWA (RRID:SCR_010910). Duplicate reads were infrequent but discarded.

ChIP-seq read density values were normalized per million mapped reads. High-confidence ChIP-seq peaks were called by MACS2 (https://github.com/taoliu/MACS) with the broad peak calling for H3K27me3, narrow peak for the rest proteins. Peaks from ChIP-seq of H3K27ac, H3K4me1, H3K4me3, H3K27me3, MYCN, WDR5 and G9a were selected based on p-value (p<10^-5^ for G9a, p<10^-7^ for the rest proteins). HOMER (RRID:SCR_010881) was used to annotate the distribution of peaks (such as enhancer, promoter, intronic, intergenic, exonic, etc.) and identify the known and *de novo* motifs.

The peak sets for MYCN, histone marks and other proteins were further analyzed using the deepTools2 suite (v3.3.0)(25). By using bamCoverage, peaks were normalized to reads per kilobase per million reads normalized read numbers (RPKM). Heatmaps and metagene plots of signal intensity of ChIP samples were generated using deepTools. Briefly, computeMatrix was used to calculate signal intensity scores per ChIP sample in a given genome region that was specified by a bed file. The output of computeMatrix was a matrix file of scores of two ChIP samples which was then used to generate the heatmaps using the plotHeatmap function and generate composite plot using the plotProfile function. For *k*-means clustering, the resulting matrix was *k*-means clustered and then visualized using plotHeatmap (--kmeans 5). For IGV sample track visualization, RPKM normalized coverage density maps (tdf files) were generated by extending reads to the average size and counting the number of reads mapped to each 25 bp window using igvtools (26).

ComputeMatrix function of the deepTools was used to generate a matrix of signal intensity of TFs of their peak centers (± 500 bp, total 1000 bp), as intensity scores in 10 bp bins. The matrix of signal intensity was further used to calculate the accumulated signal around each peak center, which was then used as the signal intensity for each TF binding site.

To find the unique and overlapped peaks between the binding sites of MYCN and its cofactors, R package, ChIPpeakAnno was applied. The function of “findOverlapsOfPeaks” was used with the “connectedPeaks” set to “min”(27).

### Statistics

The statistical analyses used throughout this paper are specified in the appropriate results paragraphs and Methods sections. Additional statistical analyses were performed using standard two-tailed Student’s *t*-test, one-way ANOVA, and the software GraphPad Prism (RRID:SCR_002798).

## Results

### MYCN governs a malignant NB cell identity by activating canonical MYC target genes and suppressing neuronal differentiation genes

*MYCN* is a well-known oncogenic driver in NB, but the molecular mechanisms by which MYCN stimulates oncogenesis remain incompletely understood. We systematically investigated *MYCN* biological functions and transcriptional activity in several NB cell lines through both loss and gain of function studies. As previously reported (19,20), the knockdown of *MYCN* in IMR32 cell line (*siMYCN_2* and *siMYCN_4*) and in KCNR, LAN5 and BE(2)C (*siMYCN_2*) resulted in a decrease of cell proliferation and increase of neurite extension (Fig. 1A-D and Fig. S1A-J). For gain of *MYCN* function studies, the non-tumorigenic SHEP NB cell line that does not express *MYCN* (*MYCN* non-amplified) was used. Overexpression of *MYCN* in SHEP for 2 days resulted in a cell morphology change with a flatter, more round cell bodies compared to control cells (Fig. S1K,L). Consistent with a previous report (28), an anchorage-independent cell proliferation assay showed that overexpression of *MYCN* in SHEP cells increased soft agar colony formation (Fig. S1M,N).

**Fig. 1.**
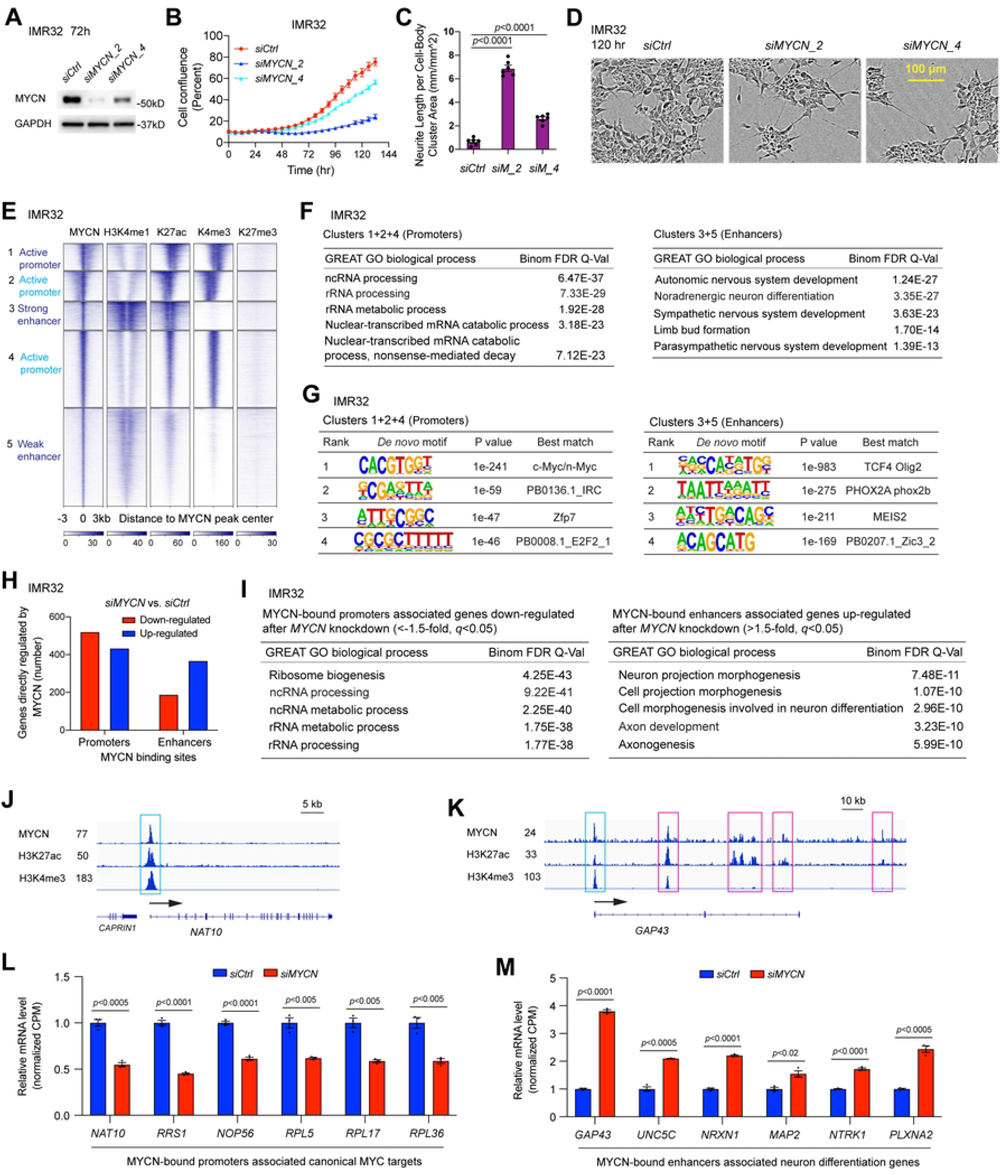
MYCN governs a malignant NB cell identity by directly activating canonical MYC target genes and suppressing neuronal differentiation genes. (**A**) The knockdown of *MYCN* using two different siRNAs in IMR32 cells for 72 h results in a decrease of MYCN at the protein levels detected by western blot assay. (**B**) The knockdown of *MYCN* using two different siRNAs in IMR32 cells results in a decrease of cell number detected by IncuCyte cell confluence assay. (**C**) and (**D**) The knockdown of *MYCN* in IMR32 cells results in an increase of neurite length shown by both the IncuCyte neurite analysis assays and phase-contrast images. (**E**) *k*-Means clustering of MYCN and histone marks ChIP-seq around MYCN binding sites of NB cell line IMR32 (±3 kb) shows that MYCN binds to proximal regulatory elements containing active promoters that are marked by H3K27ac and H3K4me3 signals, and distal regulatory elements containing strong enhancers that are marked by strong H3K4me1 and H3K27ac signals, and weak enhancers that are marked by weak H3K4me1 and H3K27ac signals. (**F**) GREAT Gene Ontology (GO) analysis indicates that MYCN-bound promoters associated genes are enriched in RNA processing and MYCN-bound enhancers associated genes are enriched in nervous system development. (**G**) HOMER motif analysis shows the enrichment of canonical E-box in the promoters and the enrichment of non-canonical E-box in the enhancers. (**H**) The combination of RNA-seq and ChIP-seq data analysis in IMR32 cells shows that more MYCN-bound promoters associated genes are down-regulated (521 vs. 434) but more MYCN-bound enhancers associated genes are up-regulated (368 vs. 189) after the silencing of *MYCN*. (**I**) GREAT GO analysis indicates that MYCN-bound promoters associated genes down-regulated after the knockdown of *MYCN* in IMR32 cells are enriched in ribosome biogenesis and RNA processing (left panel), and MYCN-bound enhancers associated genes up-regulated after the knockdown of *MYCN* are enriched in neuronal differentiation (right panel). (**J**) Signal tracks show the MYCN, H3K27ac and H3K4me3 ChIP-seq signals at the promoter of *NAT10* gene (cyan box). (**K**) Signal tracks show the MYCN, H3K27ac and H3K4me3 ChIP-seq signals at the promoter (cyan box) and enhancers (pink box) of *GAP43* gene. (**L**) The silencing of *MYCN* in IMR32 cells results in a significant down-regulation of genes involved in ribosome formation based on the RNA-seq results, and MYCN binds to the promoters of these genes. The *p*-value indicated is calculated in one-way ANOVA. CPM: counts per million. (**I**) The silencing of *MYCN* in IMR32 cells results in a significant up-regulation of genes involved in neuron projection morphogenesis based on the RNA-seq results, and MYCN binds to the enhancers of these genes. The *p*-value indicated is calculated in one-way ANOVA. CPM: counts per million.

To identify genes regulated by MYCN and understand how MYCN governs a malignant NB tumor phenotype, we performed RNA-seq analysis after silencing *MYCN* in the models described above and identified genes regulated by MYCN (Table S1). Gene set enrichment analysis (GSEA) (29,30) showed that the silencing of *MYCN* using two different siRNAs (*siMYCN_2* or *siMYCN_4*) in IMR32 resulted in a similar negative enrichment of MYC targets and ribosome biogenesis genes, whereas neuron markers and genes that positively regulate synaptic transmission were positively enriched (Fig. S1O,P). Similar results were observed when *MYCN* was knocked down in other NB cell lines by using *siMYCN_2* (Fig. S1Q,R), or when MYCN was overexpressed in SHEP cells (Fig. S1S). The loss and gain of *MYCN* function studies in multiple NB cell lines indicate that MYCN activates canonical *MYC* target genes and represses neuronal differentiation genes to govern a malignant NB cell identity.

### Genome-wide mapping of MYCN binding

To identify genes directly regulated by MYCN and the chromatin status associated with these genes, we performed ChIP-seq experiment using an MYCN antibody and antibodies that recognize different histone marks in IMR32 cells. High-confidence ChIP-seq peaks were called by MACS2 and peaks were normalized to reads per kilobase per million reads normalized read numbers (RPKM, see Materials and Methods for details). The MYCN ChIP-seq heatmap represented genome-wide stringent sets of MYCN peaks (MYCN binding sites) within the whole genome (Fig. 1E). These MYCN peaks were segmented based on their co-localizations with the histone marks (Fig. 1E) through *k*-means clustering: a total of 5 distinct clusters of MYCN peaks were identified to co-localize with differing histone marks. In general, H3K4me1 marks both active and poised enhancers, H3K27ac marks both active enhancers and promoters, H3K4me3 marks active promoters, while H3K27me3 marks repressed chromatin. Based on the peak distribution and the signal intensity of the histone marks, clusters 1, 2 and 4 of the heatmap represented active promoters or proximal regulatory regions, cluster 3 represented strong enhancers, and cluster 5 represented weak or paused enhancers based on weak H3K4me1 and H3K27ac signals (Fig. 1E). The heatmap showed that MYCN overlapped with active histone marks but not the repressive histone mark H3K27me3 (Fig. 1E). Of note, in clusters 1 and 2, we observed a shift of MYCN peaks from MYCN peak centers either to the right or to the left, which was accompanied by the shift of H3K27ac and H3K4me3 peaks, further indicating the association of MYCN binding and histone modification within these genomic loci. For all the MYCN peaks, approximately 50% are within promoters and 50% are within enhancers. Genomic regions enrichment of annotation tool (GREAT) (31) was used to analyze the peak distribution of each of these clusters. Consistent with being located in promoter and enhancer regions, the results showed that the majority of peaks for clusters 1, 2 and 4 were within 5 kb of the transcription start sites (TSS), whereas the majority of peaks for clusters 3 and 5 were 5 kb away from the TSS (data not shown). GREAT gene ontology (GO) analysis of MYCN binding sites associated genes showed that MYCN-bound clusters 1, 2 and 4 associated genes are enriched in RNA processing and regulation of gene expression, while MYCN-bound clusters 3 and 5 associated genes are enriched in nervous system development (Fig. 1F, Fig. S1T). HOMER motif analysis showed that in IMR32 cells, canonical E-box (CACGTG) was enriched in MYCN-bound promoters, whereas non-canonical E-box (CATATG) was enriched in MYCN-bound enhancers (Fig. 1G).

We further dissected MYCN binding sites and their associated genes using a publicly available MYCN ChIP-seq dataset generated in another *MYCN*-amplified cell line BE(2)C (GSE94822). Here we simply separated MYCN binding sites into two groups, which include promoter regions (-1 kb – +100 bp from TSS) and distal regulatory regions (the regions outside of the promoter) as annotated by the HOMER tool. GREAT GO analysis of MYCN binding sites associated genes showed that MYCN-bound promoter-associated genes were enriched in canonical MYC target genes that regulate RNA processing and ribosome biogenesis, whereas MYCN-bound distal regulatory regions associated genes were enriched in nervous system development (Fig. S1U).

Altogether, the ChIP-seq analyses in both IMR32 and BE(2)C cells indicate that MYCN-bound promoters are associated with canonical MYC target genes and MYCN-bound distal regulatory regions are associated with neuronal genes in NB.

### MYCN binds promoters to activate canonical MYC targets but binds to both enhancers and promoters to repress neuronal differentiation genes in NB

To investigate MYCN’s transcriptional effects on the genes it binds, we performed an integrative analysis of the MYCN ChIP-seq and RNA-seq data in IMR32 cells. For MYCN-bound promoter (defined as shown in Fig. 1E, clusters 1, 2 and 4) associated genes, a similar number of genes were down-regulated (521 genes, 54.6%) or up-regulated (434 genes, 45.4%) after the silencing of *MYCN,* while for MYCN-bound enhancer (defined as shown in Fig. 1E, clusters 3 and 5) associated genes, more genes were up-regulated (368 genes, 66.1%) than genes were down-regulated (189 genes, 33.9%) (Fig. 1H). MYCN-bound promoter-associated MYCN-activated genes (genes that were down-regulated after the silencing of *MYCN*) were significantly enriched in ribosome biogenesis and RNA processing (Fig. 1I, left panel), whereas MYCN-bound promoter-associated, MYCN-repressed genes (genes that were up-regulated after the silencing of *MYCN*) were enriched in pons development and response to axon injury (Fig. S1V). The MYCN-bound enhancer-associated MYCN-repressed genes were significantly enriched in neuronal differentiation (Fig. 1I, right panel), whereas MYCN-bound enhancer-associated MYCN-activated genes were enriched in chordate embryonic development (Fig. S1W). RPKM normalized signal tracks showed a MYCN ChIP-seq peak at the promoter of *NAT10* (N-Acetyltransferase 10), a gene required for ribosome biogenesis (Fig. 1J). For the neuronal differentiation gene, *GAP43* (growth associated protein 43), a single MYCN ChIP-seq peak was observed at the promoter while multiple peaks were observed at its enhancers (Fig. 1K). We found that in addition to binding to the enhancers of neuronal genes in NB cells, MYCN also binds to the promoters of these genes. The expression changes of representative MYCN-bound promoter-associated ribosome biogenesis genes after MYCN depletion were shown in Fig. 1L, and expression changes of representative MYCN-bound enhancer-associated neuronal genes were shown in Fig. 1M. These results indicate that MYCN activates canonical MYC target genes mainly through binding to promoters in NB, whereas MYCN binds to both enhancers and promoters to suppress neuronal differentiation genes.

### MYCN binds to promoters to activate canonical MYC targets but binds to enhancers to repress skeletal muscle genes in rhabdomyosarcoma

To investigate whether the mechanism by which MYCN regulates gene expression in NB is consistent in other embryonal tumor marked by MYCN overexpression, we interrogated publicly available ChIP-seq data from rhabdomyosarcoma (RMS) cell line RH4 that expresses high levels of *MYCN* (GSE83728) (32). *K*-means clustering results showed that in RH4 cells, approximately 25% of MYCN peaks were within promoters while approximately 75% of MYCN peaks were within enhancers (Fig. 2A). In RH4 cells, MYCN bound to both active promoters and active enhancers but not to repressed chromatin indicated by different histone marks (Fig. 2A). MYCN-bound promoter-associated genes were enriched in canonical MYC target genes such as genes that regulate RNA processing (Fig. 2B, left panel), whereas MYCN-bound enhancer-associated genes were enriched in skeletal system development (Fig. 2B, right panel). Silencing of *MYCN* in RH4 cells resulted in a decrease in cell proliferation (Fig. S2A-C). GSEA of the RNA-seq data (Table S1) showed that the knockdown of *MYCN* in RH4 cells resulted in a negative enrichment of canonical MYC target genes that are involved in RNA processing and ribosome biogenesis, as well as a positive enrichment of myogenic differentiation signature genes (HSMM_UP) (33) (Fig. 2C). To investigate the association of MYCN binding sites and their target gene expression, we further performed an integrative analysis of MYCN ChIP-seq and RNA-seq data derived from RH4 cells. Similar to our observation in NB cells, we found that more of MYCN-bound enhancer-associated genes were up-regulated (221 vs. 58) after *MYCN* silencing in RMS cells (Fig. 2D). GO analysis indicated that MYCN-bound promoter-associated genes that were down-regulated after the silencing of *MYCN* in RH4 cells were enriched in RNA processing (Fig. 2E, left panel), whereas MYCN-bound enhancer-associated genes that were up-regulated after the silencing of *MYCN* were enriched in muscle system processes (Fig. 2E, right panel). Neither MYCN-bound promoter-associated MYCN-repressed genes nor MYCN-bound enhancer-associated MYCN-activated genes showed significantly enriched biological processes. Signal tracks showed a MYCN ChIP-seq peak at the promoter of *NAT10* gene that is required for ribosome biogenesis (Fig. 2F), while MYCN ChIP-seq peaks were observed both at the promoter and enhancers of muscle gene, *MYL4* (myosin light chain 4) (Fig. 2G). Representative MYCN-bound promoter-associated MYCN-activated genes that are involved in ribosome biogenesis were shown in Fig. 2H, and representative MYCN-bound enhancer-associated MYCN-repressed genes that are involved in muscle system processes were shown in Fig. 2I.

**Fig. 2.**
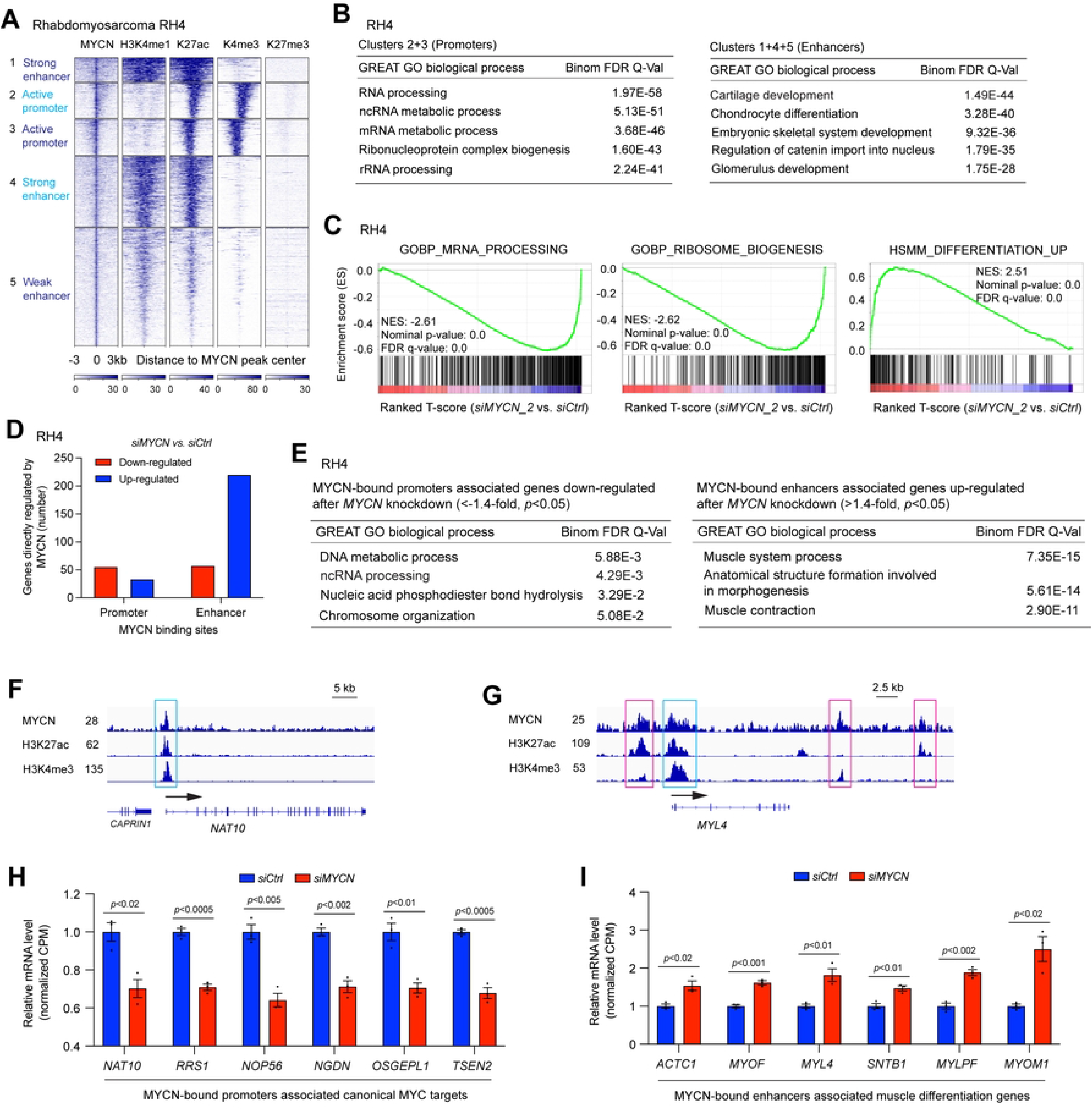
MYCN binds to the promoters to activate canonical MYC targets and binds to the enhancers to repress muscle differentiation genes in RMS. (**A**) *k*-Means clustering of MYCN and histone marks ChIP-seq around MYCN binding sites of rhabdomyosarcoma cell line RH4 (±3 kb). (**B**) GREAT GO analysis indicates that MYCN-bound promoters associated genes are enriched in RNA processing and MYCN-bound enhancers associated genes are enriched in cartilage system and skeletal system development. (**C**) GSEA of the RNA-seq data shows that the knockdown of *MYCN* in RH4 cells for 72 h results in a negative enrichment of canonical MYC target genes that are involved in RNA processing and ribosome biogenesis, as well as a positive enrichment of human myogenic differentiation signature genes (HSMM_UP). (**D**) The combination of RNA-seq and ChIP-seq data analysis in RH4 cells shows that more MYCN-bound promoters associated genes are down-regulated (56 vs. 34) but more MYCN-bound enhancers associated genes are up-regulated (221 vs. 58). (**E**) GREAT GO analysis indicates that MYCN-bound promoters associated genes down-regulated after the knockdown of *MYCN* in RH4 cells are enriched in RNA processing (left panel), and MYCN-bound enhancers associated genes up-regulated after the knockdown of *MYCN* are enriched in muscular system process (right panel). (**F**) Signal tracks show the MYCN, H3K27ac and H3K4me3 ChIP-seq signals at the promoter of *NAT10* gene (cyan box). (**G**) Signal tracks show the MYCN, H3K27ac and H3K4me3 ChIP-seq signals at the promoter (cyan box) and enhancers (pink box) of *MYL4* gene. (**H**) The silencing of *MYCN* in RH4 cells results in a significant down-regulation of genes involved in ribosome formation based on the RNA-seq results, and MYCN binds to the promoters of these genes. The *p*-value indicated is calculated in one-way ANOVA. CPM: counts per million. (**I**) The silencing of *MYCN* in RH4 cells results in a significant up-regulation of genes involved in muscle system processes based on the RNA-seq results, with MYCN binding to the enhancers of these genes. The *p*-value indicated is calculated in one-way ANOVA. CPM: counts per million.

Thus, in two distinct pediatric cancers, NB and RMS, our results demonstrate that MYCN directly activates canonical MYC target genes mainly through binding to the promoters of these genes, while repressing tissue-specific differentiation genes mainly through binding to both enhancers and promoters of these genes (Fig. 1I-M, Fig. 2E-I).

### *MYCN* depletion alters histone modifications on its target genes

We next asked whether the activation of canonical MYC target genes and repression of neuronal differentiation genes by MYCN are associated with changes of histone modifications after MYCN depletion by ChIP-seq analysis. To compare the ChIP-seq signal intensity in control and *MYCN* silenced samples, the ChIP-seq peaks were RPKM normalized. While changes in MYCN levels led to decreases in the average MYCN ChIP-seq signals, there were no changes in the average ChIP-seq signals for histone marks (Fig S3A). Analysis of MYCN binding focused on TSS showed similar results (Fig. S3B).

Furthermore, we focused on genes whose expression was modulated by changes in MYCN expression and directly bound by MYCN. When silencing *MYCN*, the integrative analysis of the MYCN ChIP-seq and RNA-seq merged data identified 601 genes directly activated by MYCN (bound by 936 MYCN peaks) whose expression decreased with *MYCN* silencing, and 625 genes directly suppressed by MYCN (bound by 1420 MYCN peaks) whose expression increased with *MYCN* silencing. Ingenuity Pathway Analysis (IPA) showed that MYCN-bound genes whose expression decreased after the silencing of *MYCN* were enriched in RNA post-transcriptional modification and protein synthesis (Fig. 3A), whereas the MYCN-bound genes that were up-regulated after *MYCN* silencing were positively enriched in neuronal differentiation (Fig. 3B).

**Fig. 3.**
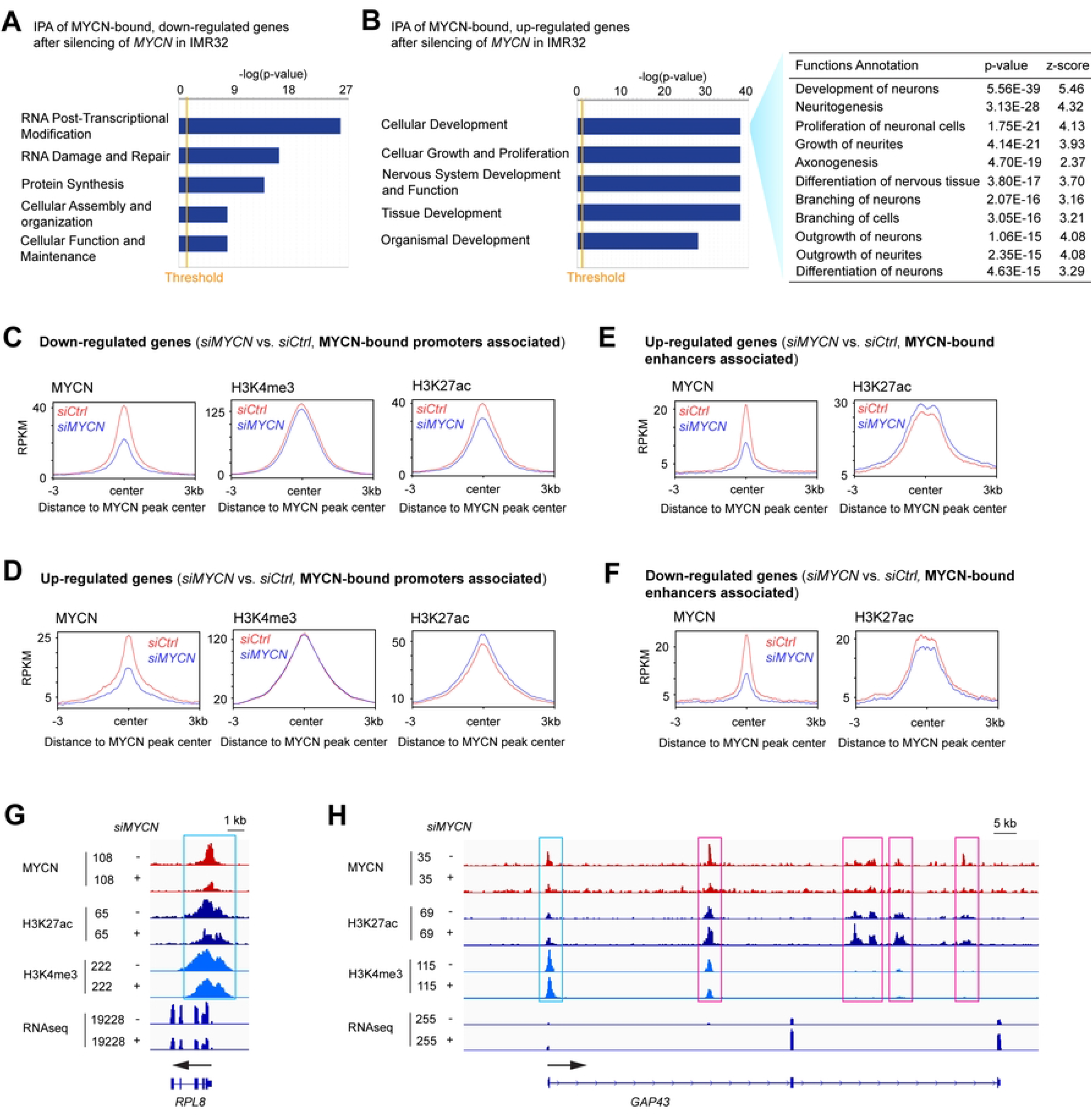
MYCN depletion alters histone modifications on its target genes. (**A**) The integrative analysis of the MYCN ChIP-seq and RNA-seq merged data by ingenuity pathway analysis (IPA) shows that the MYCN-bound, down-regulated genes after the silencing of *MYCN* in IMR32 cells are enriched in RNA post-transcriptional modification and protein synthesis. (**B**) IPA of the RNA-seq data shows that the MYCN-bound up-regulated genes after the silencing of *MYCN* are positively enriched in neuronal differentiation. (**C**) When focuses on MYCN-bound promoters associated, down-regulated genes, metagene plots show that the silencing of *MYCN* results in a decrease in the average ChIP-seq signals of H3K4me3 and H3K27ac at the MYCN peak center. (**D**) When focusing on MYCN-bound promoters associated, up-regulated genes, metagene plots show that the silencing of *MYCN* results in an increase in the average ChIP-seq signal of H3K27ac at the MYCN peak center. (**E**) When focusing on MYCN-bound enhancers associated, up-regulated genes, metagene plots show that the silencing of *MYCN* results in an increase in the average ChIP-seq signal of H3K27ac at the MYCN peak center. (**F**) When focusing on MYCN-bound enhancers associated, down-regulated genes, metagene plots show that the silencing of *MYCN* results in a decrease in the average ChIP-seq signal of H3K27ac at the MYCN peak center. (**G**) Signal tracks show decreases in MYCN, H3K27ac and H3K4me3 ChIP-seq signals at the promoter of *RPL8* after the depletion of MYCN (cyan box). (**H**) Signal tracks show increases in H3K27ac ChIP-seq signals at the promoter (cyan box) and enhancers (pink boxes) of *GAP43* after the depletion of *MYCN*.

Since we discovered that MYCN binds to the promoters of MYCN activated canonical MYC target genes and binds to the enhancers of MYCN repressed neuronal genes (Fig. 1I), we next focused on how the silencing of *MYCN* affects the epigenetic modifications at MYCN binding sites within these promoters and enhancers. By focusing on the promoters of MYCN-bound down-regulated genes due to the silencing of *MYCN*, we found a decrease in the average ChIP-seq signals of active promoter marks H3K4me3 and H3K27ac at the MYCN peak center after the silencing of *MYCN* (Fig. 3C). When focused on the promoters of MYCN-bound, up-regulated genes after *MYCN* silencing, we found an increase of the average ChIP-seq signal of H3K27ac at the MYCN peak center (Fig. 3D). By focusing on the enhancers of MYCN-bound genes after *MYCN* silencing, we found an increase in the average H3K27ac ChIP-seq signal at the MYCN peak center for up-regulated genes (Fig. 3E) but for down-regulated genes there was a decrease in the H3K27ac signal (Fig. 3F). For example, signal tracks showed decreases of MYCN, H3K27ac and H3K4me3 ChIP-seq signals at the promoter of *RPL8* (ribosomal protein L8) after the depletion of MYCN (Fig. 3G), whereas signal tracks for the neuronal differentiation gene, *GAP43*, showed decreases of MYCN ChIP-seq signals and increases of H3K27ac ChIP-seq signals at both the promoter and enhancers after the depletion of MYCN (Fig. 3H). These results suggest that MYCN directly activates canonical MYC target genes through increasing promoter activity, whereas MYCN directly represses neuronal differentiation genes through repressing enhancer activity.

### Interactome assay to identify MYCN novel cofactors

TFs recruit cofactors to remodel the chromatin and/or modify histones to regulate gene transcription. To identify cofactors that mediate the transcriptional activity of MYCN on activating canonical MYC target genes and repressing differentiation genes, we performed co-immunoprecipitation (co-IP) coupled with mass-spectrometry assessments using two different MYCN antibodies to identify protein interactors of endogenous MYCN in IMR32 cells (Fig. S4A, Fig. 4A). Each MYCN antibody pulled down around 400 protein partners, of which 337 were overlapped (Table S2). The analysis confirmed multiple known MYCN and MYC partners, such as MAX, TRRAP, topoisomerases IIA and IIB (TOP2A and TOP2B) (34–36). Ingenuity pathway analysis of MYCN protein partners showed that 224 of the 337 MYCN protein partners (62%) are nuclear proteins (Fig. S4B, Table S2). The DAVID tool (37) was used to annotate MYCN protein partners within the nucleus and showed that these proteins were enriched in different categories such as chromosome organization, chromatin organization, intracellular ribonucleoprotein complex, DNA replication and mRNA splicing (Fig. S4C). Importantly, in support of our discovery, when compared to the recently identified, high-confidence interactors for c-MYC (Table S2) that were characterized in Hela cells based on proximity-dependent biotinylation technique (BioID) (35), we found that 89 of the 337 MYCN interactors also interact with c-MYC (Fig. 4A, Table S2).

**Fig. 4.**
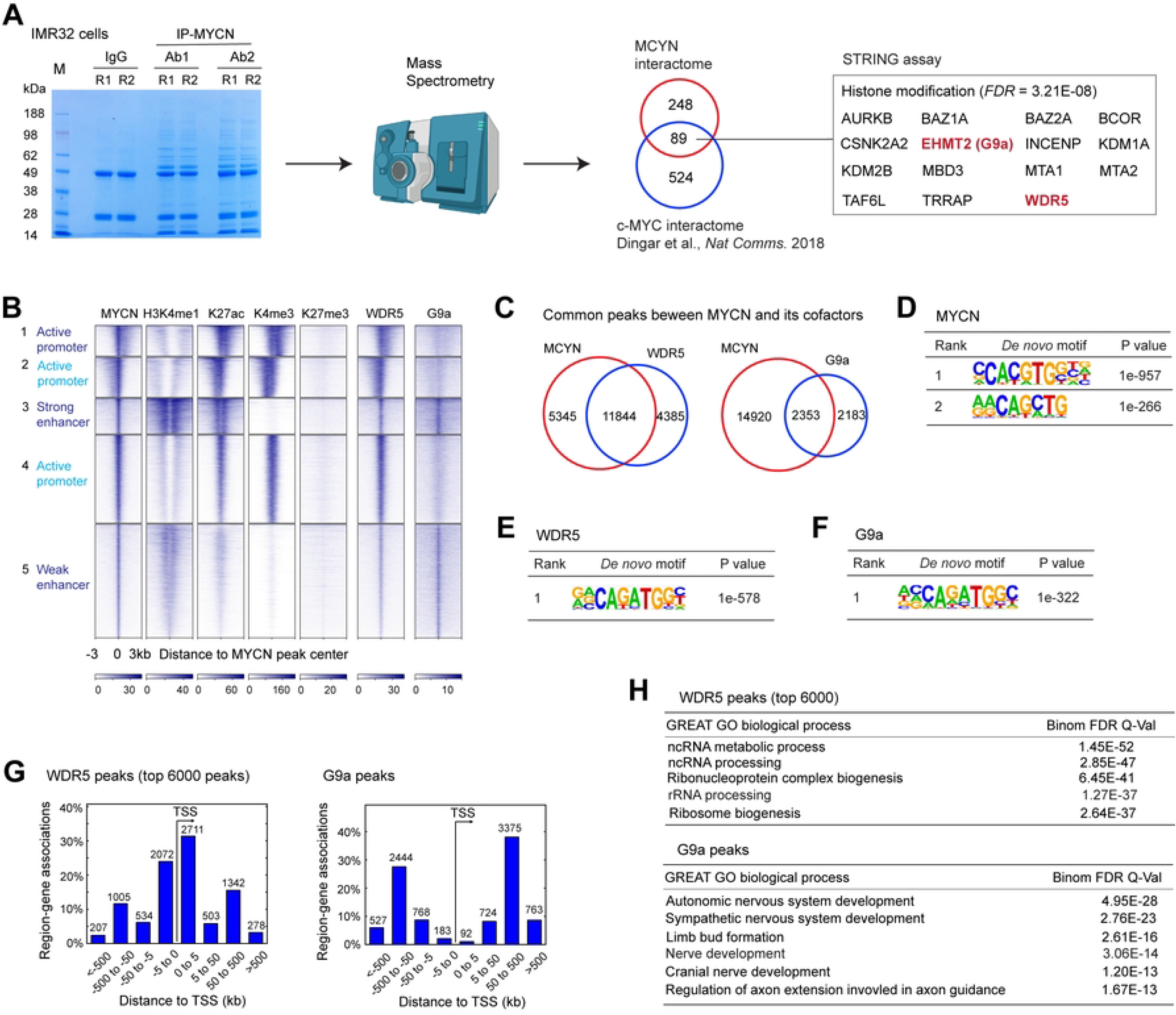
Genome-wide co-localization of MYCN and its cofactors. (**A**) Protein bands identified in the MYCN pulldown products in the protein gel (stained by Coomassie blue) are used for in-gel digestion and mass spectrometry sequencing (left panel). Discovered MYCN protein partners are compared with published c-MYC interactors (middle panel). Gene ontology analysis of the common protein interactors using STRING tool identifies protein enriched in histone modification (right panel). (**B**) *K*-Means clustering of MYCN and histone marks ChIP-seq, MYCN cofactors ChIP-seq and CRC TFs ChIP-seq around MYCN binding sites of NB cell line IMR32 (±3 kb) shows the overlapping binding sites between MYCN and its cofactors. (**C**) ChIPPeakAnno analysis shows the number of common and unique peaks between MYCN and its cofactors. (**D**-**F**) HOMER motif analysis shows that both canonical and non-canonical E-box are enriched in MYCN peaks, and non-canonical E-box is enriched in WDR5 and G9a peaks. (**G**) GREAT peak distribution analysis shows that around 50% of WDR5 peaks are within 5kb from the TSS (left panel), while only less than 5% of G9a peaks are within 5 kb from the TSS (right panel). (**H**) GREAT GO analysis indicates that WDR5-bound top ranked peaks associated genes are enriched in RNA processing and ribosome biogenesis (top panel), while G9a-bound peaks associated genes are enriched in nervous system development (bottom panel).

Next, we prioritized those MYCN interactors that are involved in histone modifications and those considered promising targets for anti-cancer therapies. Gene ontology analysis of the 89 proteins that interact with both MYCN and c-MYC using the STRING protein-protein interaction tool (38) identified 15 proteins that are significantly enriched in histone modifications (Fig. 4A). Among these proteins, we were particularly interested in histone WDR5 (a component of histone methylation complex) and G9a (histone methyltransferase, also known as EHMT2). WDR5 was found to be a coactivator of c-MYC and could be a focal point for anti-c-MYC therapies (39,40). In NB, WDR5 was found to form a protein complex with MYCN at the MDM2 promoter to regulate MDM2 transcription (11), but whether MYCN cooperates with WDR5 to regulate global gene transcription in NB has not been characterized. G9a was found to be a corepressor of c-MYC (41). It has been shown that inhibition of G9a promotes neuronal differentiation of human bone marrow mesenchymal stem cells (42). Moreover, G9a was found to be essential in many types of cancers including NB (43). However, the association between MYCN and G9a has not been investigated. We performed MYCN co-IP and western blot analysis in IMR32 cells and confirmed that MYCN could pull down both WDR5 and G9a (Fig. S4D). Consistent with our discovery, a recent MYCN interactome assay performed in HEK293 cells with over-expressed MYCN (Table S2) (36) also revealed that WDR5 and G9a interact with MYCN.

### Genome-wide co-localization of MYCN and its cofactors

To investigate the genome-wide interactions of MYCN and the coactivator WDR5 or corepressor G9a, we performed ChIP-seq analysis of MYCN, WDR5 and G9a in IMR32 cells. Aligned to the clusters of MYCN binding sites and histone marks, we found that WDR5 bound to both promoters and enhancers, although with a stronger signal intensity at the promoter regions (Fig. 4B). In contrast, G9a predominantly bound to the enhancers (Fig. 4B). ChIPPeakAnno analysis showed that 73% WDR5 and 52% G9a binding sites overlapped with MYCN binding sites (Fig. 4C). HOMER motif scan showed that the top 2 MYCN binding motifs are canonical and non-canonical E-boxes, while the non-canonical E-box was found to be enriched in WDR5 and G9a binding sites (Fig. 4D-F). Consistent with the *k*-means clustering analysis, GREAT peak distribution analysis showed that few G9a binding sites were within 5 kb of TSS (<5%), while 55% of WDR5 binding sites were within 5 kb of TSS (Fig. 4G). The genome-wide co-localization of MYCN with either WDR5 or G9a suggested that each of these cofactors cooperates with MYCN to regulate a subset of MYCN target genes. GREAT GO analyses showed that MYCN WDR5 ChIP-seq peak-associated genes were enriched in RNA processing and ribosome biogenesis (Fig. 4H, top panel), suggesting its potential role as a MYCN coactivator. On the other hand, G9a binding site-associated genes were enriched in nervous system development (Fig. 4H, bottom panel), suggesting its potential role as a MYCN corepressor. These results suggest that MYCN cooperates with WDR5 to regulate canonical MYC target genes while MYCN cooperates with G9a to regulate neuronal differentiation genes.

### *MYCN* silencing alters the genomic DNA binding of its cofactors

To identify whether MYCN recruits its cofactors to its binding sites, we performed ChIP-seq experiments using MYCN, WDR5 and G9a antibodies in *siCtrl* and *siMYCN* transfected IMR32 cells. The silencing of *MYCN* did not alter the steady-state protein levels of G9a but caused a 50% decrease in WDR5 based on densitometric analysis (Fig. 5A). ChIP-seq results showed that the knockdown of *MYCN* in IMR32 cells caused a 15% decrease in WDR5 ChIP-seq peak numbers (18263 to 15612 peaks) (Table S3), and a 64% decrease in G9a ChIP-seq peak numbers (6723 to 2428 peaks) (Table S4).

**Fig. 5.**
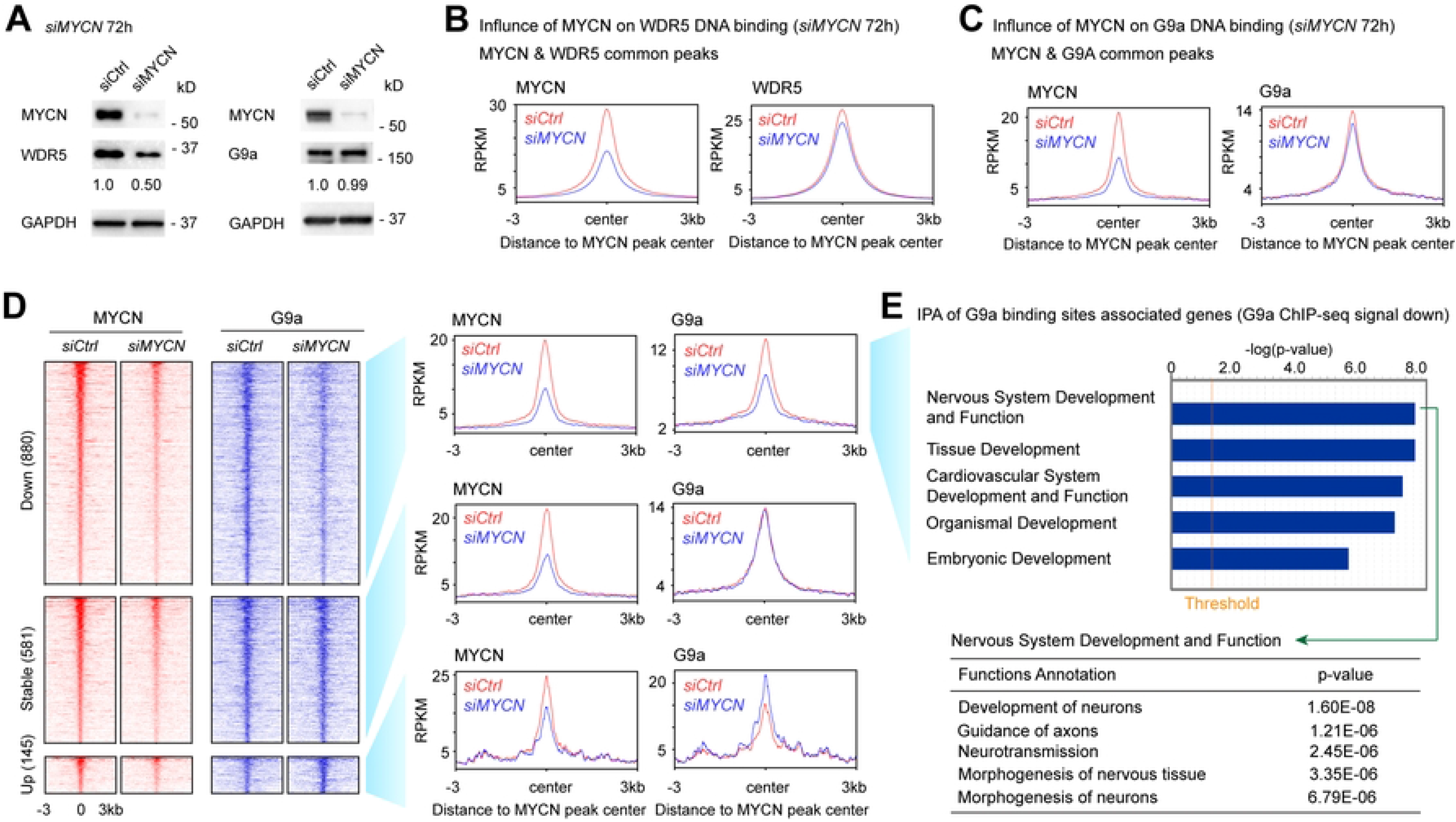
Silencing of *MYCN* selectively alters genomic DNA binding of its cofactors. (**A**) Western blot and densitometric analyses show the effect of 72 h knockdown of *MYCN* in IMR32 cells on the expression of its cofactors at the protein levels. (**B**) Metagene plots show that the knockdown of *MYCN* results in a decrease of average MYCN and WDR5 ChIP-seq signal at the MYCN peak center of the MYCN and WDR5 overlapped binding sites. (**C**) Metagene plots show that the knockdown of *MYCN* results in a decrease in average MYCN and G9a signal at the MYCN peak center of MYCN and G9a overlapped binding sites. (**D**) ChIP-seq heatmaps (left) and profiles (right) of MYCN and G9a in control and *MYCN* knockdown IMR32 cells around MYCN binding sites (±3 kb) of MYCN and G9a overlapped peaks. (**E**) IPA of the G9a ChIP-seq data shows that the G9a binding sites with decreased ChIP-seq signal (>1.2-fold decrease of G9a signal after the silencing of *MYCN*) associated genes are enriched in nervous system development.

By focusing on MYCN and WDR5 overlapped peaks, metagene plots showed a 50% decrease in the average MYCN ChIP-seq signals and a 5% decrease in the average WDR5 ChIP-seq signals at the summit of MYCN peak centers (Fig. 5B) after silencing of *MYCN*. Signal tracks for the ribosome gene *RPL8* promoter and the cell adhesion molecule coding gene *PVR* promoter showed decreases in WDR5 binding after *MYCN* silencing (Fig. S5A). The silencing of *MYCN* resulted in a 50% decrease of WDR5 at the protein level but was accompanied by only a slight decrease of WDR5 ChIP-seq peak numbers and average ChIP-seq signal intensity, suggesting that MYCN is not required for WDR5 to bind to DNA.

When evaluating MYCN and G9a overlapped peaks, metagene plots show a 5% decrease in the average G9a ChIP-seq signals at the summit of MYCN peak centers after silencing MYCN (Fig. 5C). Signal tracks for the *KCNK3* gene showed that the knockdown of *MYCN* resulted in a decrease of MYCN and G9a binding signals within the *KCNK3* intron (Fig. S5B). *MYCN* silencing resulted in a subtle decrease in the average G9a ChIP-seq signals (Fig. 5C) but dramatically reduced G9a ChIP-seq peak numbers (Table S4), thus, we further analyzed MYCN and G9a overlapping peaks by dissecting them into three clusters. The first cluster was classified as down-regulated (down) peaks, which included G9a peaks with at least > 1.2-fold decrease of G9a ChIP-seq signal after the silencing of *MYCN* (Fig. 5D); the second cluster was classified as not-altered (stable) peaks, which included G9a peaks with < 1.1-fold changes of G9a ChIP-seq signal after the silencing of *MYCN* (Fig. 5D); whereas the third cluster was classified as increased (up) peaks, which included G9a peaks with > 1.2-fold increase of G9a ChIP-seq signal after the silencing of *MYCN* (Fig. 5D). More G9a binding peaks had decreases in their ChIP-seq signal than the ones that were stable or had increased signals in *MYCN* silenced cells (Fig. 5D), which was consistent with the observation of a reduced number of G9a ChIP-seq peaks after the silencing of *MYCN* (Table S4). Ingenuity pathway analysis (IPA) of the G9a peaks associated genes with a decreased or stable ChIP-seq signal revealed that these genes were enriched in nervous system development (Fig. 5E, Fig. S5C). Furthermore, the genes associated with G9a binding sites that had increased ChIP-seq signals were enriched in the development of other tissues such as hair and skin development (Fig. S5D). Our results indicated that MYCN selectively recruits G9a to MYCN binding sites that are associated with neuronal genes.

### MYCN cofactors facilitate MYCN binding to DNA

Recent studies showed that target gene recognition by c-MYC is not solely dependent on interactions with MAX, but also depends on other proteins including WDR5 (39,40). Mutations in c-MYC that disrupt the WDR5 interaction result in a significant decrease in the binding of c-MYC to its target genes (39). To investigate whether WDR5 is required for MYCN to bind to DNA, we performed ChIP-seq analysis before and after the silencing of *WDR5*. Western blot results showed that the silencing of *WDR5* did not alter the expression of MYCN protein (Fig. 6A).

**Fig. 6.**
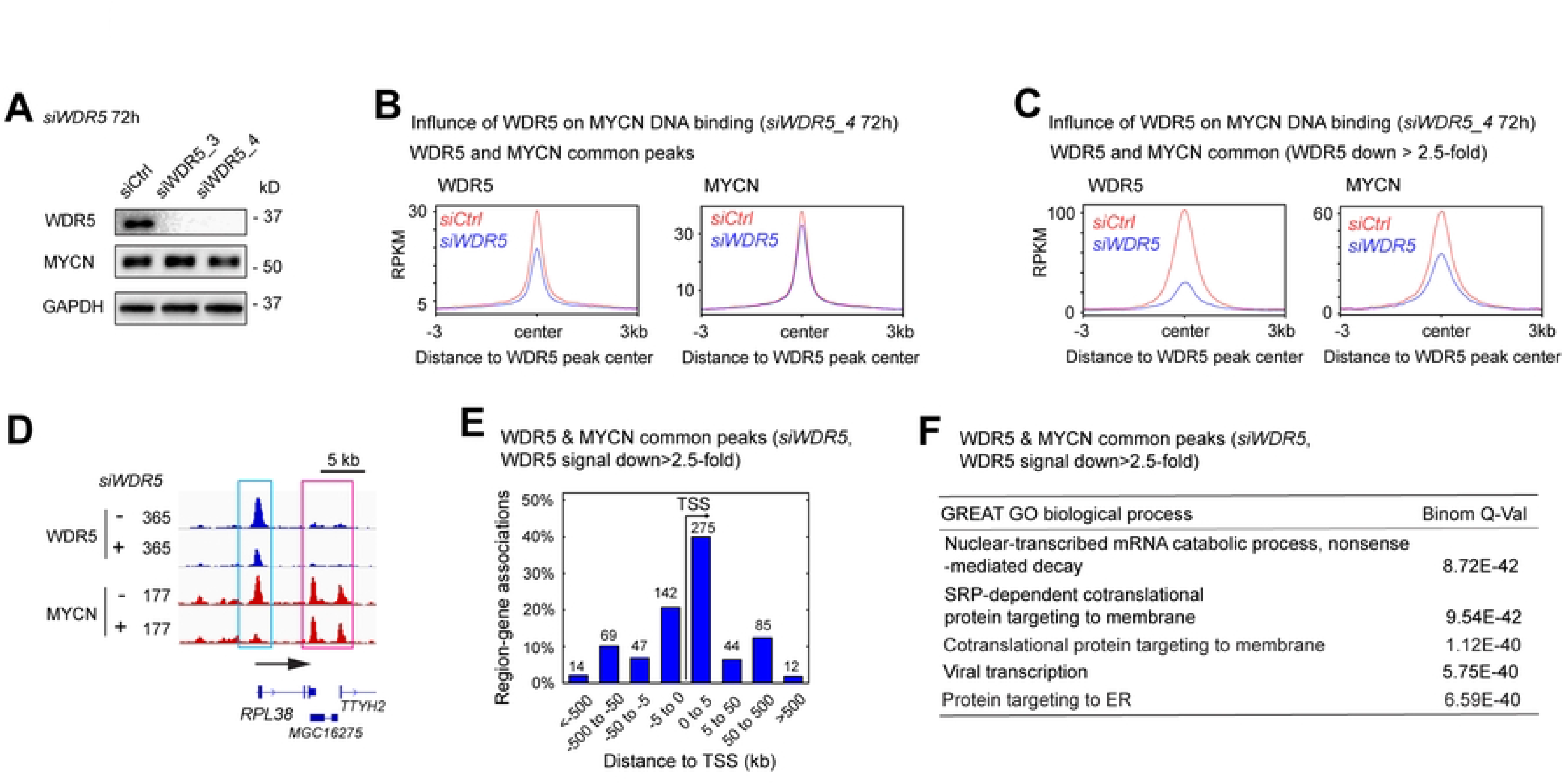
MYCN cofactors assist MYCN to bind to DNA. (**A**) Western blot analysis shows that the knockdown of *WDR5* using siRNAs for 72 h decreases in WDR5 protein but has no effect on MYCN expression. (**B**) and (**C**) Metagene plots show that the knockdown of *WDR5* results in a decrease in average WDR5 and MYCN ChIP-seq signals at the WDR5 peak center, and the decrease of average MYCN signal is more dramatic when focused on WDR5 and MYCN overlapped binding sites with >2.5-fold decrease of WDR5 ChIP-seq signal intensity. (**D**) Signal tracks show that the knockdown of *WDR5* results in a decrease of MYCN signal at the promoter of the *RPL38* gene (cyan box) but not at the *TTYH2* gene locus (pink box). (**E**) When focused on WDR5 and MYCN overlapped peaks with >2.5-fold decrease of WDR5 ChIP-seq signals, GREAT peak distribution analysis shows that around 60% of WDR5 & MYCN overlapped peaks are within 5kb away from the TSS, and (**F**) GREAT GO biological process analysis shows that these WDR5 and MYCN overlapped peaks associated genes are enriched in RNA processing and protein synthesis.

When focused on WDR5 and MYCN overlapped binding sites, metagene plots showed that the silencing of *WDR5* resulted in a decrease in the average ChIP-seq signal of MYCN at the WDR5 and MYCN overlapped binding sites (Fig. 6B). We next investigated the genomic loci where WDR5 assisted MYCN binding by focusing on MYCN-bound promoters and enhancers as defined by the histone marks (Fig. 1E). Metagene plots showed that the knockdown of *WDR5* resulted in an approximately 20% decrease in average MYCN ChIP-seq signal at the summit of MYCN-bound promoters and a 10% decrease in MYCN signal at the summit of MYCN-bound enhancers (Fig. S6A,B), suggesting that WDR5 has a greater effect on MYCN binding to promoters compared to enhancers. Of note, the knockdown of *WDR5* resulted in >90% depletion of WDR5 protein levels detected by western blot (Fig. 6A), but it only reduced the number of WDR5 ChIP-seq peaks by 52% (15448 to 7404 peaks) (Table S5), and reduced WDR5 ChIP-seq signal intensity by approximately 40% at the summit of the WDR5 peak center (Fig. 6B, left panel), suggesting that the remaining WDR5 after *WDR5* knockdown is able to bind to certain genomic loci with high WDR5 binding affinity. For example, signal tracks showed that the silencing of *WDR5* did not alter WDR5 and MYCN ChIP-seq signals at the promoter of the *NAT9* gene and *TMEM104* gene (Fig. S6C). Thus, we focused on the WDR5 and MYCN overlapped binding sites with >2.5-fold decrease of WDR5 ChIP-seq signals after the silencing of *WDR5* (Fig. 6C, left panel), to investigate how the depletion of WDR5 influenced MYCN-DNA interaction on these binding sites. This time the metagene plots showed an obvious decrease (50%) in MYCN ChIP-seq signals at these genomic loci (Fig. 6C, right panel), indicating that WDR5 is required for MYCN to bind DNA on these genomic loci. As an example, signal tracks showed that the silencing of *WDR5* resulted in a >70% decrease of both WDR5 and MYCN ChIP-seq signals at the promoter of ribosome biogenesis gene *RPL38* (Fig. 6D, cyan box).

To investigate which genes were cooperatively bound by WDR5 and MYCN, we focused on the WDR5 and MYCN overlapped binding sites with >2.5-fold decrease in WDR5 ChIP-seq signals after the silencing of *WDR5*. GREAT peak distribution analysis showed that 60% of these peaks were within 5 kb from the TSS, while the remaining were 5 kb away from the TSS (Fig. 6E). This indicated that WDR5 mainly assisted MYCN binding to promoters. GREAT GO analysis indicated that these WDR5 and MYCN overlapped peak-associated genes are significantly enriched in protein synthesis, RNA processing and ribosome biogenesis, but not in nervous system development (Fig. 6F, Table S6). Taken together, these findings indicate that WDR5 mainly assists MYCN to bind to the promoters that are associated with MYCN activated canonical MYC targets.

### The depletion of MYCN cofactors antagonizes MYCN-mediated gene expression changes

Our study indicated that WDR5 and G9a interact with MYCN and that these proteins colocalize to the promoters or enhancers of genes they regulate (Fig. 4D). To investigate whether the presence of WDR5 or G9a is necessary to regulate MYCN target genes, we silenced *WDR5*, *G9a* or *MYCN* using siRNAs for 48 h in IMR32 cells and performed RNA-seq analysis. Western blot results showed that the silencing of *WDR5* or *G9a* had no effect on MYCN expression at this time point (Fig. S7A). Like the results of silencing of *MYCN* for 72 h (Fig. S1O), GSEA results showed that the 48 h silencing of *MYCN* in IMR32 cells resulted in a significant negative enrichment of canonical *MYC* target genes involved in ribosome biogenesis, RNA processing, ribosome formation and cytoplasmic translation (Fig. S7B), and resulted in a significant positive enrichment of neuronal genes involved in axon development, neuron differentiation, glutamatergic synapse and neuron projection guidance (Fig. S7C). To compare genes regulated by MYCN, WDR5, and G9a in IMR32 cells after 48 h silencing, we generated “MYCN activated canonical MYC targets” gene sets by defining genes that were significantly down-regulated after the silencing of *MYCN* in the gene sets of ribosome biogenesis, RNA processing, ribosome formation and cytoplasmic translation (Table S7). In parallel, we generated “MYCN repressed neuronal genes” gene sets by defining those genes that were significantly up-regulated after the silencing of *MYCN* in the gene sets of axon development, neuron differentiation, glutamatergic synapse, positive regulation of synaptic transmission and neuron projection guidance (Table S7).

The silencing of *WDR5* resulted in a significant negative enrichment of “MYCN activated canonical MYC targets” gene sets such as genes involved in ribosome formation and cytoplasmic translation (Fig. 7A). GSEA results did not exhibit significant positive enrichment of neuronal genes after the silencing of *WDR5*. The RNA-seq results were consistent with the discovery that WDR5 and MYCN overlapped peaks-associated genes were enriched in canonical MYC targets, but not enriched in the nervous system development (Fig. 6F, Table S6).

**Fig. 7.**
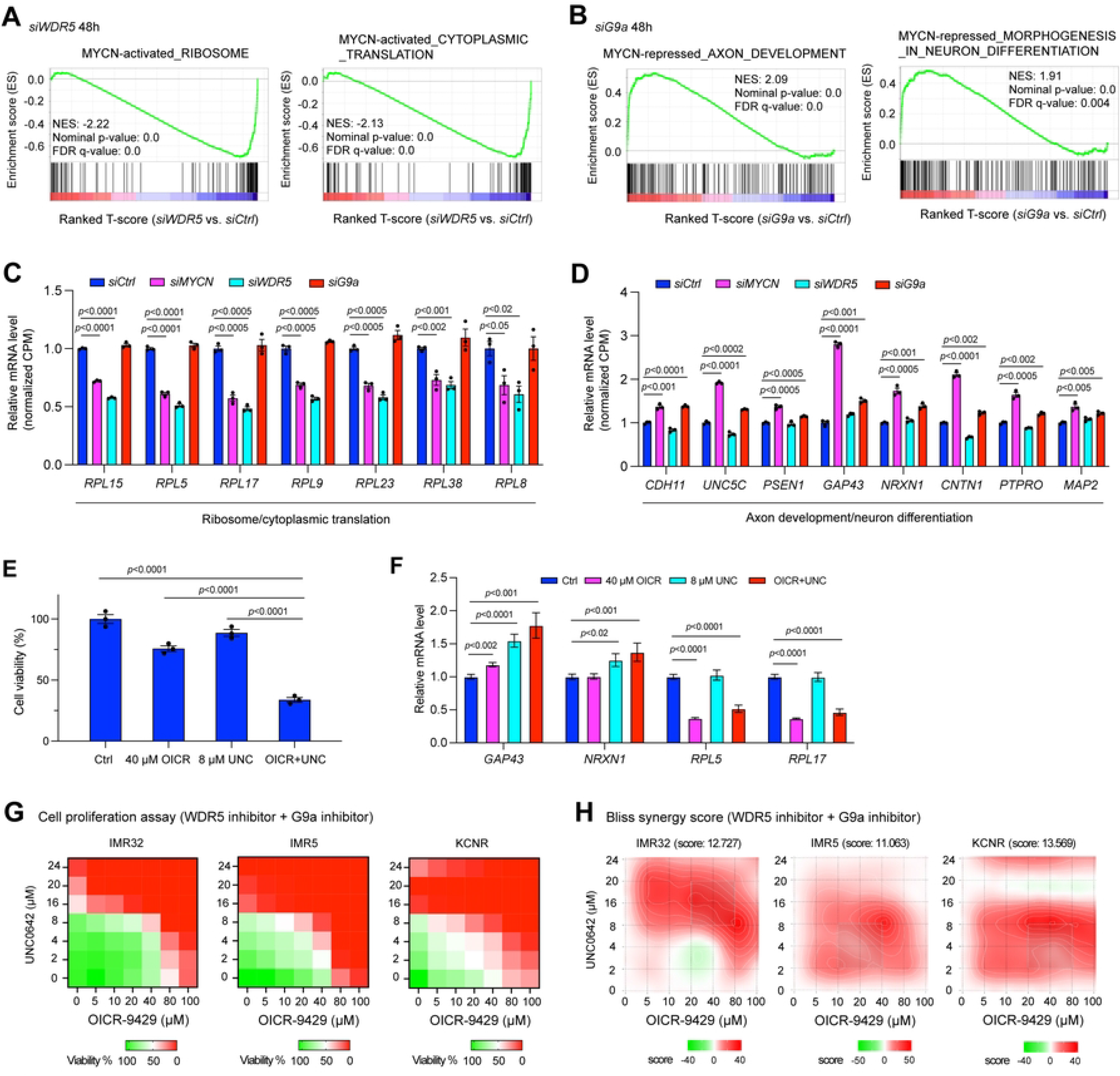
The depletion of MYCN cofactors antagonizes MYCN-mediated gene transcription regulation, which makes them potential therapeutic targets. (**A**) GSEA shows that the silencing of *WDR5* results in a significant negative enrichment of genes involved in ribosome formation and protein synthesis that are activated by MYCN. (**B**) GSEA shows that the silencing of *G9a* results in a significant positive enrichment of genes involved in axon development and neuron differentiation that are repressed by MYCN. (**C**) The silencing of either *MYCN* or *WDR5* but not *G9a* results in a significant down-regulation of genes involved in ribosome formation and protein translation based on the RNA-seq results. The *p*-value indicated is calculated in one-way ANOVA. CPM: counts per million. (**D**) The silencing of either *MYCN* or *G9a* but not *WDR5* results in a significant up-regulation of genes involved in axon development and neuron differentiation based on the RNA-seq results. The *p*-value indicated is calculated in one-way ANOVA. CPM: counts per million. (**E**) CellTiter-Glo assay shows the drug effect of the OICR-9429 (OICR, 40 µM) + UNC0642 (UNC, 8 µM) treatment on IMR32 cell viability at 72 h. (**F**) Realtime PCR shows that the inhibition of both WDR5 and G9a results in a significant down-regulation of ribosomal genes and up-regulation of neuronal genes. (**G**) Heatmaps show the percentage of cell viability after different doses of WDR5 inhibitor OICR-9429 and G9a inhibitor UNC0642 treatment in *MYCN*-amplified NB cell lines. Cells are treated with the drugs for 72 h and cell viability is measured by CellTiter-Glo Cell Viability Assay. (**H**) SynergyFinder online tool is used for bliss synergistic analysis to evaluate the synergistic effect of the combination treatment in *MYCN*-amplified NB cell lines shown in (**G**).

G9a predominantly binds to the enhancers (Fig. 4B) and G9a-bound peak-associated genes were enriched in nervous system development (Fig. 4H). GSEA results showed that the silencing of *G9a* resulted in a significant positive enrichment of “MYCN repressed neuronal genes” gene sets such as genes involved in axon development, neuron differentiation, glutamatergic synapse, and neuron projection guidance (Fig. 7B, Fig. S7D), while no significant negative enrichment of canonical MYC target genes was observed after the silencing of *G9a*.

Representative genes commonly down-regulated after the silencing of *MYCN* or *WDR5* that belong to the gene sets of “MYCN activated canonical MYC targets” were shown in Fig. 7C. The same group of genes were not down-regulated after the silencing of *G9a* (Fig. 7C). Representative genes commonly up-regulated after the silencing of *MYCN* or *G9a* that belong to the gene sets of “MYCN repressed neuronal genes” were shown in Fig. 7D. The same group of genes was not up-regulated after the silencing of *WDR5* (Fig. 7D). These results indicated that the silencing of *WDR5* antagonizes MYCN-mediated activation of canonical MYC target genes, while the silencing of *G9a* antagonizes MYCN-mediated repression of neuronal genes.

### Targeting both MYCN coactivator and corepressor simultaneously

MYCN is an oncogenic driver but is still considered undruggable directly by small molecules due to the flexibility of its protein structure. Thus, it has been proposed to target the druggable MYCN cofactors with enzymatic activity. Indeed, it has been shown that the treatment of NB cells with WDR5 inhibitor suppressed cell proliferation (11,44), and G9a inhibitor treatment suppressed NB growth (45,46). Our preceding studies showed that WDR5 cooperates with MYCN to activate canonical MYC target genes while G9a cooperates with MYCN to repress neuronal differentiation genes, and the silencing of either *WDR5* or *G9a* only affects the expression of a subset of MYCN target genes. Therefore, we hypothesized that a more efficient strategy to drug MYCN activities would be to target these cofactors simultaneously.

First, we evaluated whether the genetic inhibition of *WDR5* or *G9a* affects NB cell proliferation. DepMap (https://depmap.org/portal/) CRISPR library screen data analysis showed that *WDR5* was essential in all the NB cell lines while half the NB cells were partially dependent on *G9a* to survive or proliferate based on the CRISPR dependence score (Fig. S7E). Of note, the *WDR5* or *G9a* dependency was not dependent on *MYCN* amplification status (Fig. S7E), which is possible due to the high expression of *c-Myc* in *MYCN* non-amplified NB cell lines (47), while WDR5 and G9a have been shown to mediate c-MYC function in the other types of cancers (39,41). Consistent with the DepMap CRISPR library screen results, we found that the genetic silencing of *WDR5* or *G9a* using siRNAs in IMR32 cells resulted in a decrease in NB cell proliferation (Fig. S7F), supporting the dependency of NB cells on each of these cofactors.

Next, as a proof of concept, we evaluated the ability of a small molecule inhibitor selective for WDR5 and one for G9a on MYCN target genes’ expression and NB cell proliferation. OICR-9429 is a WDR5 WIN site inhibitor that displaces WDR5 from chromatin (48). UNC0642 is a G9a catalytic inhibitor and across breast cancer cell lines, the anti-proliferative response to UNC0642 correlates with MYC sensitivity and gene signatures (41). Representative IMR32 cell proliferation assay showed that while using a single dose of OICR-9429 or UNC0642 that only slightly reduced cell viability (10-25%), the combination of these two drugs dramatically reduced cell viability (>60%) after 72 h drug treatment (Fig. 7E), which indicated a potential for a synergistic effect. Using the same dose of OICR-9429 or UNC0642 to treat IMR32 cells as used in the cell proliferation assay (Fig. 7E), realtime quantitative PCR results showed that only the inhibition of WDR5 significantly decreased the expression of ribosome gene *RPL5* and *RPL17*, and that only the inhibition of G9a significantly increased expression of neuronal gene *GAP43* and *NRXN1*, whereas the combination of WDR5 and G9a inhibitors’ treatment resulted in both an upregulation of neuronal gene *GAP43* and *NRXN1*, and repression of the ribosomal genes *RPL5* and *RPL17* (Fig. 7F). Further cell proliferation assays showed that the treatment of NB cells with increased dosages of either OICR-9429 or UNC0642 alone reduced the number of viable NB cells (Fig. 7G), while the combination of OICR-9429 and UNC0642 synergistically reduced NB cell proliferation with an average bliss synergy score greater than 10 in all tested *MYCN*-amplified NB cell lines (Fig. 7G,H). These results indicate that inhibiting both MYCN coactivators and corepressors is necessary to repress both the active and repressive activity of MYCN, thereby resulting in a synergistic effect on suppressing NB cell proliferation.

## Discussion

MYCN is a common oncogene in many types of cancers, yet how MYCN regulates global gene expression has not been well-characterized. This is necessary if one wants to explore an indirect approach to targeting MYCN for therapeutic purposes. Through genome-wide and proteomic approaches, we have identified critical interactions between MYCN and the transcriptional coactivator WDR5 and corepressor G9a, and mapped their interactions at a genome level in NB cells. We find that WDR5 facilitates MYCN binding to genomic DNA to activate canonical MYC target genes involved in protein synthesis, which occurs mainly via promoter binding. In contrast, MYCN recruits corepressor G9a to bind enhancers and this functions to directly repress neuronal differentiation gene expression (Fig. 8). MYCN requires these cooperative interactions to mediate its oncogenic transcriptional program by using a coactivator to stimulate growth supporting pathways and a corepressor to inhibit the potential growth inhibiting pathways that would naturally occur as cells differentiate (Fig. 8). In this way MYCN orchestrates global gene expression and governs the malignant NB cell identify. Targeting the cofactors that mediate these two pathways antagonizes the dysregulated MYCN activity and more effectively suppresses NB tumor cell growth.

**Fig. 8.**
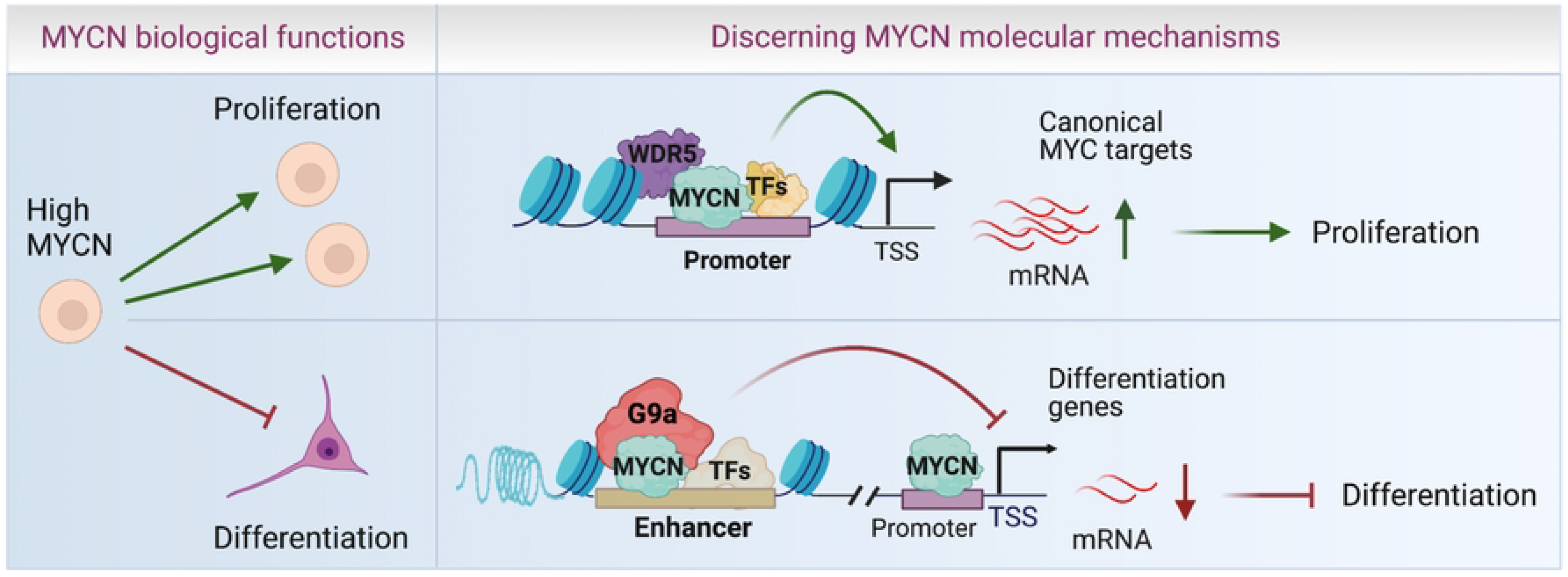
Schematic diagram of MYCN action. WDR5 assists MYCN to bind promoters and up-regulate canonical MYC target genes to stimulate cell proliferation, whereas MYCN recruits G9a to enhancers to down-regulate neuronal differentiation genes and inhibit cell differentiation.

Although it is known that MYCN activates canonical MYC targets involved in ribosome biogenesis, protein synthesis and RNA processing (1,6,7), here we find at a genome-wide level that this is mainly through MYCN binding to promoters. MYCN has been reported to repress neuronal differentiation genes (8), but whether this was direct or indirect had only been assessed on a small number of genes. Here, we demonstrate that in NB cells, MYCN binds to both the promoters and the enhancers of neuronal differentiation genes, and the depletion of *MYCN* from these binding sites results in a significant up-regulation of these genes. Previous studies showed that MYCN cooperates with MIZ1 or SP1 to repress gene transcription through binding to the MIZ1 or SP1 binding site within the promoters (49,50). For the first time we find that in addition to promoter binding, MYCN utilizes enhancer binding to repress gene expression in NB cells, as the depletion of MYCN from these enhancers is associated with activated histone modifications as shown by the increase in H3K27ac ChIP-seq signals. This observation is consistent with the well-known concept that enhancers control cell-type-specific gene expression (51). Moreover, our discovery of the repression of neuronal differentiation genes by MYCN is consistent with the observation that the silencing of *MYCN* in NB cells results in an increase in neuronal differentiation. In another MYCN driven tumor, RMS, MYCN binds to enhancers and directly represses muscle differentiation genes, suggesting a general mechanism by which enhancer-bound MYCN represses differentiation genes. Our model in which MYCN selectively activates canonical MYC targets through binding promoters and represses cell lineage specific differentiation genes through binding enhancers more fully rationalizes the different oncogenic cellular effects driven by MYCN in different types of cancer cells.

Our study discerns the molecular mechanisms of how MYCN activates canonical MYC target genes and represses neuronal genes. The coactivator WDR5 has been shown to interact with c-Myc and MYCN (11,39). WDR5 was found to recruit c-MYC to chromatin and regulate the expression of protein synthesis genes (39,40). Consistent with this, we find that WDR5 facilitates MYCN genome binding to activate canonical MYC target genes involved in ribosome biogenesis and protein synthesis. We demonstrate that many neuronal differentiation genes are repressed by MYCN and MYCN binds to the enhancers of these genes. MYCN has been shown to alter bivalent epigenetic marks (H3K4me3 and H3K27me3) by recruiting PRC2 complex to transcriptionally repress expression of CLU gene (14). Components of the PRC2 complex have not been identified in our MYCN interactome assay, and MYCN did not colocalize with H3K27me3 on the genome (Fig. 1E), suggesting that the direct repression of most of the neuronal genes by MYCN is not via PRC2 recruitment. It is possible that MYCN regulates some of these neuronal genes indirectly through PRC2 since it is known that MYCN activates EZH2 expression (52). A recent study indicated that G9a cooperates with c-MYC to repress gene transcription (41). G9a is responsible for H3K9 dimethylation that is known to be associated with transcriptional repression (53,54). In our study, we find that MYCN recruits G9a to the enhancers of neuronal differentiation genes, which possibly catalyzes H3K9me2, establishing a repressive chromatin environment at MYCN binding sites to decrease enhancer activity. A limitation of our study is that we have not been able to successfully perform a H3K9me2 ChIP-seq to determine how the recruitment of G9a to MYCN binding sites affects H3K9me2 status despite the use of most commercially available H3K9me2 antibodies. Nevertheless, our findings that MYCN co-localizes with G9a on enhancers associated with neuronal differentiation genes, and the finding that the depletion of *G9a* antagonizes MYCN-mediated repression of neuronal genes, support a model in which G9a functions as a MYCN corepressor of MYCN to suppress these cell-type specific differentiation genes. As others have proposed (55), it is highly likely MYCN engages in several protein complexes rather than forming a single defined complex in regulating gene transcription. We find that MYCN mainly cooperates with WDR5 at the promoters while cooperating with G9a at the enhancers, suggesting that MYCN forms different protein complexes with WDR5 and G9a. In addition to G9a, our MYCN interactome assay identified other corepressors or chromatin remodeling complexes such as the NuRD complex, suggesting that these corepressors might also participate with MYCN to contribute to the suppression of certain neuronal differentiation genes.

In tumors driven by MYCN, therapeutic targeting of MYCN has been a long-sought goal, but has remained challenging due to its structure flexibility (5). The intrinsic enzyme activity of cofactors offers the potential for developing strategies aimed at the indirect therapeutic targeting of MYCN. It has been shown that a WDR5 inhibitor was used to treat NB when WDR5 was found to be a coactivator of MYCN (11,44). In our study, we demonstrate that WDR5 only mediates the active transcriptional activity of MYCN in activating protein synthesis genes and RNA processing genes. Thus, targeting WDR5 alone does not fully suppress the transcriptional activity of MYCN. We find that G9a mediates the repressive transcriptional activity of MYCN in repressing neuronal differentiation genes in NB, and this led to a strategy that simultaneous inhibition of both WDR5 and G9a would more fully target MYCN oncogenic transcriptional activities. Indeed, the combination of WDR5 inhibitor and G9a inhibitor synergistically suppressed NB cell proliferation. This discovery highlights that targeting both coactivators and corepressors of an oncogenic TF simultaneously enables more precise therapy.

In summary, genome-wide mapping of MYCN binding and transcriptome analysis indicates that MYCN bind to promoters to activate canonical *MYC* target genes, whereas MYCN binds both enhancers and promoters to repress tissue specific differentiation genes. Our results indicate that the oncogenic competence of MYCN is mediated by a combination of its coactivators including WDR5, and its corepressors including G9a. MYCN forms a dynamic complex system with different cofactors on different genomic loci to control the chromatin landscape, guide the expression of genes, and determine the malignant cancer cell identity. The targeting of WDR5 and G9a simultaneously antagonizes MYCN-mediated gene regulation to synergistically suppress NB cell proliferation. The understanding of the mechanistic underpinnings of MYCN oncogenic activity provides a rationale to target both these cofactors simultaneously, which has important therapeutic implications for patients with *MYCN-*driven tumors.

## DATA AVAILABILITY

All the home generated RNA-seq and ChIP-seq can be found in the Gene Expression Omnibus (GEO) database. GEO accession number for data generated in this study is GSE208424. RNA-seq of *MYCN* silencing for 72 h in IMR32 and LAN5 can be found under GEO accession number GSE183641 (56). GEO accession number for publicly available ChIP-seq data derived from BE2C cells is GSE94822. GEO accession number for publicly available ChIP-seq data of MYCN, and histone marks derived from rhabdomyosarcoma cell line RH4 is GSE83728.

## SUPPLEMENTARY DATA

Supplementary data include one single PDF file that includes 7 supplementary figures and 8 supplementary tables in MS Excel.

## ACKNOWLEDGEMENTS

We thank Dr. John Glod for the critical review of the manuscript. We are grateful for the Colgate University Program that supported Amanda Ciardiello work at NCI. We thank Bao Tran, Jyoti Shetty and Yongmei Zhao from NCI Sequencing Facility for DNA and RNA sequencing. We thank Ming Zhou, Thorkell Andresson from NCI Protein Characterization Laboratory core facility for mass-spec assay. This work utilized the computational resources of the NIH HPC Biowulf cluster (http://hpc.nih.gov). The schematic diagram was created with BioRender.com.

## AUTHOR CONTRIBUTIIONS

Z.L. and C.J.T designed the study, wrote the manuscript, and coordinated the entire study. Z.L., X.Z., M.X., JH and AC performed the experiments. Z.L., X.Z. and H.L. performed bioinformatic analyses. J.F.S. reviewed ChIP-seq and bioinformatic analyses. All the authors have reviewed the manuscript.

